# Multi-omics of a rice population identifies genes and genomic regions in rice that bestow low glycemic index and high protein content

**DOI:** 10.1101/2024.03.21.586029

**Authors:** Saurabh Badoni, Erstelle Pasion-Uy, Sakshi Kor, SungRyul Kim, Gopal Misra, Rhowell Tiozon, Reuben James Q. Buenafe, Ana Rose Ramos-Castrosanto, Vipin Pratap, Inez Slamet-Loedin, Julia von Steimker, Saleh Alseekh, Alisdair R. Fernie, Ajay Kohli, Gurudev S. Khush, Nese Sreenivasulu

**Author notes:** contributed equally (Saurabh Badoni, Erstelle Pasion-Uy).

## Abstract

To address the growing incidences of increased diabetes and to meet the daily protein requirements, we developed low glycemic index (GI) rice varieties with protein yield exceeding 14%. In the development of recombinant inbred lines using Samba Mahsuri and IR36 amylose extender as parental lines, we identified quantitative trait loci (QTLs) and genes associated with low GI, high amylose content (AC), and high protein content (PC). By integrating genetic techniques with classification models, this comprehensive approach identified candidate genes on chromosome 2 (qGI2.1/qAC2.1 spanning the region from 18.62Mb to 19.95Mb), exerting influence on low GI and high amylose. Notably, the phenotypic variant with high value was associated with the recessive allele of the starch branching enzyme 2b (sbeIIb). The genome-edited sbeIIb line confirmed low GI phenotype in milled rice grains. Further, combinations of alleles from the highly significant SNPs from the targeted associations and epistatically interacting genes showed ultra-low GI phenotypes with high amylose and high protein. Metabolomics analysis of rice with varying AC, PC, and GI revealed that the superior lines of high AC and PC, and low GI were preferentially enriched in glycolytic and amino acid metabolism, whereas the inferior lines of low AC and PC and high GI were enriched with fatty acid metabolism. The high amylose high protein RIL (HAHP_101) was enriched in essential amino acids like lysine. Such lines may be highly relevant for food product development to address diabetes and malnutrition.

**Significance Statement:** The increasing global incidence of diabetes calls for the development of diabetic friendly healthier rice. In this study, we developed recombinant inbred rice lines with milled rice exhibiting ultra-low to low glycemic index and high protein content from the cross between Samba Mahsuri and IR36 amylose extender. We performed comprehensive genomics and metabolomics complemented with modeling analyses emphasizing the importance of *OsSbeIIb* along with additional candidate genes whose variations allowed us to produce target rice lines with lower glycemic index and high protein content in a high-yielding background. These lines represent an important breeding resource to address food and nutritional security.

## Introduction

Globally, 537 million adults are presently suffering from diabetes and it is estimated that this will reach up to 783 million by 2045 (1). Among these incidences, type 2 diabetes accounts for 90-95% of diabetes (1). Low- and middle-income countries contribute to more than 75% of diabetes incidences, with Asia accounting for 60% of the global diabetic population (1), wherein 90% of the polished rice is produced and consumed in Asia. Rice, being a staple for more than half of the global population, is consumed primarily in the polished form which is loaded with easily digestible starch (of total ∼90% starch; dry weight). This means most cultivated rice varieties exhibit a high glycemic index (GI) (2). Increased consumption of high GI food and highly processed products combined with a sedentary lifestyle collectively leads to an increased risk of type 2 diabetes and other non-communicable diseases (2). A significant number of individuals in developing countries depend heavily on staple foods such as rice for their daily calorie and protein requirements (3). The identification of rice varieties characterized by low GI with enhanced levels of protein content (particularly those high in essential amino acids; EAAs) is imperative to improve public health among low and middle-income communities in Asia.

High resistant starch (RS)/low-glutelin *sbeIIb/Lgc1* lines were generated by CRISPR-Cas9-induced site-specific mutations in *SBEIIb* in an elite low-glutelin japonica rice cultivar (4). According to this study, the mutant lines showed increases of about two times in amylose content (AC) and 6% in RS with low glutelin content which is the primary rice seed storage protein. These findings point to the trade-off between increased amylose and grain protein content. Furthermore, the storage protein content of the grain also has an impact on slow digestibility, leading to low GI (5). Among the cereal grains, rice generally has the lowest protein content, but its net protein utilization is the highest (6). Only a small number of rice protein content genes have been identified to date. Thus, it is of pivotal importance to understand the genetic factors to achieve a pyramided strategy of high amylose combined with high protein lines in a high-yielding background to lower the GI and create healthier protein supplements.

Herein, we used QTL-seq analysis using bulk segregants analysis (BSA-Seq) combined with next-generation sequencing of two pools, the high amylose and high protein (HAHP) lines, and the low amylose and low protein (LALP) lines. QTL and targeted association studies of F_6_ recombinant inbred lines (RILs) were used to identify the genes responsible for regulating HAHP. We further unraveled the metabolomic signatures of these lines also determined the protein content and essential amino acid (EAA) composition of the pure protein isolate. The functional validation of the *sbeIIb* allele by gene editing confirmed the low GI phenotype. We contend that the candidate genes and pathways unraveled through this investigation leading to rice breeding lines with ultra-low GI, high AC, and PC can be deployed to develop diabetic friendly gluten-free rice protein supplements with enriched essential amino acids for human health.

### Genetic regions influencing high amylose and high protein in milled rice grains identified by bulk segregant analysis (BSA-Seq)

A broad variation in amylose content (AC, 18.1 to 36.8%) was observed in the F_3_ population developed from the cross of IR36 amylose extender mutant (*IR36ae*) with high amylose and the recipient line Samba Mahsuri with superior grain quality (Figure S1). The *in vitro* GI values for the high and low AC bulk pools were 46 (ultra-low GI) and 65 (intermediate GI), while the grain protein content (PC) ranged from 6% to 15.5%. Five significant QTL peaks [*qseqAC1*.*1* (6.02-6.18 Mb), *qseqAC2*.*1* (7.74-8.61 Mb), *qseqAC2*.*2* (17.15-19.95 Mb), *qseqAC2*.*3* (20.72-21.90 Mb), and *qseqAC6*.*1* (28.83-29.67 Mb)] from Chromosomes 1, 2, and 6 were detected influencing AC based on BSA-seq analysis (Figure 1A, Table S1.1). The *qseqAC2*.*1* region showed high and moderate effect SNPs (from candidate genes involved in storage protein development (LOC_Os02g15070 – *OsEnS-32*/Glutelin precursor B6), grain size and shape (LOC_Os02g14730 – *LARGE GRAIN1*) and cell wall structure (LOC_Os02g14900 – 1,3-beta-glucan synthase), as well transcription factors including OsEnS-34/PROLAMIN BOX BINDING FACTOR (LOC_Os02g15350) (Table S2.1). A highly significant QTL, *qseqAC2*.*2* (17.15-19.95 Mb), also confirmed in the QTL analysis of F_6_ population, comprised numerous candidate genes involved in carbohydrate, amino acid metabolism, protein synthesis, and lipid metabolism including *OsSBEIIb* (LOC_Os02g32660), which is a known major gene modulating starch branching in rice grain and two other candidate genes regulating sucrose degradation (LOC_Os02g32730 – *OsNIN3*/*neutral/alkaline invertase 3*, and LOC_Os02g33110 – *OsCIN1/Cell-wall invertase 1*), along with genes encoding germin-like glycoproteins, ankyrin repeat family proteins, and amino acid activation (Table S1.1, S2.1). The *qseqAC2*.*3* also showed significant candidate genes for sucrose degradation (LOC_Os02g34560 *OsNIN8*/*neutral/alkaline invertase 8*), elongation of α-glucan chains (DP12-30) during amylopectin synthesis (LOC_Os02g51070 – *OsSSIIb*/*Soluble starch synthase IIb*), and grain length regulation (LOC_Os02g51320 - *Positive Regulator of Grain Length 2*) (Table S1.1, S2.1).

**Figure 1.**
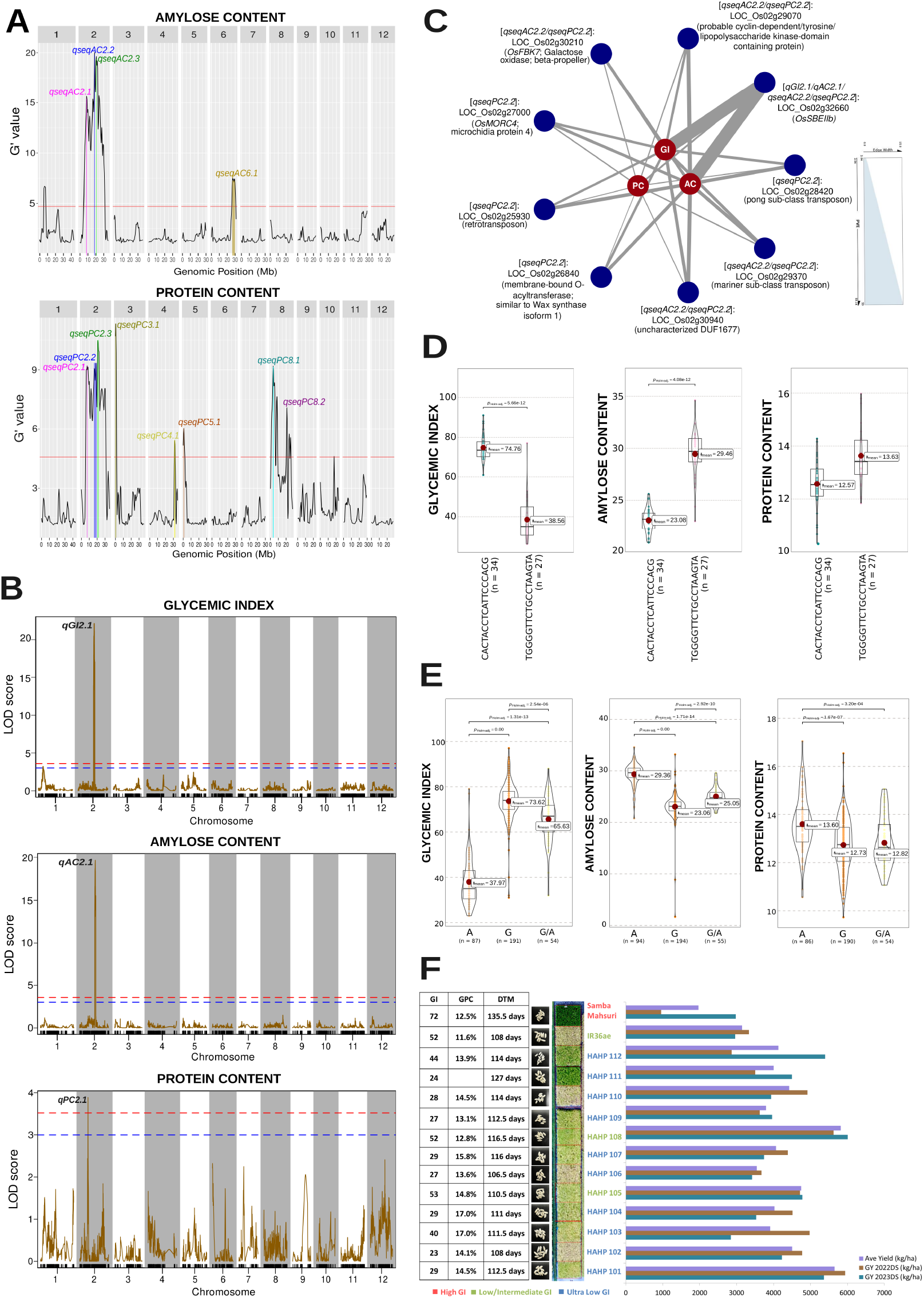
Summary of candidate QTLs and genes showing significant phenotypic variations for GI, AC, and PC of recombinant inbred lines based on various genomics approaches. (A) Significant QTL peaks associated with AC and PC based on bulk-segregant analysis on pooled contrasting F2-derived F3 recombinant inbred lines (RILs). (B) Significant QTL peaks associated with GI, AC, and PC based on composite interval mapping of F5-derived F6 RILs. (C) Candidate genes from QTL regions filtered after targeted association analysis showing phenotypic variations for GI, AC, and PC. (D) Combinations of alleles from nine candidate genes including OsSBEIIb showed a range of GI, AC, and PC properties which can be utilized for table consumption or further product development. (E) Phenotypic distributions of F5-derived F6 RILs grouped based on a single SNP change in OsSBEIIb. (F) GI, PC, days-to-maturity (DTM), and grain yield for two different years (2022 and 2023 dry season) of selected ultra-low and low GI RILs for further breeding advancement and product development.

For grain protein content, seven QTLs [(*qseqPC2*.*1* (7.06-8.44 Mb), *qseqPC2*.*2* (15.01-20.20 Mb), *qseqPC2*.*3* (20.71-21.37 Mb), *qseqPC3*.*1* (0.41-0.95 Mb), *qseqPC4*.*1* (31.69 Mb), *qseqPC5*.*1* (0.86-1.0 Mb), *qseqPC8*.*1* (3.50-4.06 Mb) *and qseqPC8*.*2* (20.65-21.31 Mb)] were identified (Figure 1A, Table S1.2). The QTL *qseqPC2*.*1* overlapping *qseqAC2*.*1* consists of *Glutelin B6* (LOC_Os02g15070), *Glutelin D* (LOC_Os02g15090), and *Glutelin 7* (LOC_Os02g15150), as well as some genes involved in sugar transport (LOC_Os02g13560 - *Tonoplast monosaccharide transporter 2*) and meiotic recombination during fertilization (LOC_Os02g13810 - *Human enhancer of invasion 10*) (Table S1.2, S2.2). On the other hand, *qseqPC2*.*2* (overlapping with *qseqAC2*.*2* and *qAC2*.*1*) contained the genes including *OsSBEIIb, OsNIN3*, and *OsCIN1* as well as candidate genes encoding glutelin (LOC_Os02g25860 – *Glutelin A*), *TRANSPARENT TESTA GLABRA 1B* (LOC_Os02g32430), fasciclin-like arabinogalactan proteins, germin-like glycoproteins, and ankyrin repeat family/tetratricopeptide domain-containing proteins, along with enzymes such as glycosyltransferases, aminotransferase, galacturonosyltransferase, nucleoside-diphosphate-sugar epimerase, digalactosyldiacylglycerol synthase, and metallo-beta-lactamase-trihelix chimera (Table S1.2). Almost completely overlapping with *qseqAC2*.*3, qseqPC2*.*3* also comprised *OsNIN8* and *PLANT GLYCOGENIN-LIKE STARCH INITIATION PROTEIN A3* (LOC_Os02g35020) with multiple high effect variants for PC (Table S2.2).

### Genetic regions influencing ultra-low GI, high amylose, and high protein confirmed through QTL analysis and targeted association studies from the F_5_-derived F_6_ population

QTL analysis of the F_6_ population revealed two significant QTLs: *qGI2*.*1*/*qAC2*.*1* (spanning the region from 18.62Mb to 19.95Mb influencing ultra-low GI and high amylose phenotypes) and *qPC2*.*1* (interval of 10.13Mb to 10.33Mb influencing grain protein %) (Figure 1B, Table S3). The fine-mapped *qGI2*.*1*/*qAC2*.*1* genetic region overlapping with *qseqAC2*.*2* confirmed the locus identified through BSA-seq, comprising numerous candidate genes with high-effect variants significantly associated with GI and AC. The region overlapped *OsSBEIIb* and confirmed important candidate genes such as *OsCIN1* (Table S1, S2). On the other hand, *qPC2*.*1* showed additional candidate genes such as LOC_Os02g17620 (isochorismatase) associated with PC.

Targeted association analysis revealed that important candidate genes identified from the BSA-seq analysis from the RIL population showed a total of 215 SNPs from 134 candidate genes significantly associated with AC and PC (Table S4.1). A single nucleotide change to homozygous A allele in *SBEIIb* contributed 57.2% PVE to GI, 60% PVE to AC and 8% PVE to PC (Figure 1C, 1D). Some candidate genes affecting AC and PC as well as GI and are highly expressed in early developing seeds include those encoding galactose oxidase/beta propeller (OsFBK7: LOC_Os02g30210), probable lipopolysaccharide kinase domain-containing protein (LOC_Os02g29070), DUF1677 (LOC_Os02g30940), membrane-bound O-acyltransferase (LOC_Os02g26840; similar to Wax synthase isoform 1), OsMORC4 (LOC_Os02g27000), and some transposons (LOC_Os02g25930, LOC_Os02g28420, LOC_Os02g29370) (Figure 1C). The allelic combinations of the significant SNPs from these candidate genes revealed distinct GI classes ranging from ultra-low, intermediate, to high GI with specific groups having high to medium AC and PC. The superior set of lines exhibited ultra-low GI (38.56 mean), with high AC (29.46 mean), and high PC (13.63 mean), whilst the inferior set of lines exhibited high GI 79.00 mean), intermediate AC (25.17 mean), and intermediate PC (11.78 mean).

Other critical candidate genes identified for AC and PC were LOC_Os02g15070 (*Glutelin B6/OsEnS-32*), LOC_Os02g33110 (*OsCIN1*), LOC_Os02g34560 (*OsNIN8*), LOC_Os02g31290 (LARGE1 controlling grain size and weight), LOC_Os02g34630 (MYB transcription factor), LOC_Os02g28970 (microspore and tapetum regulator 1), LOC_Os02g33230 (nucleoside-diphosphate-sugar epimerase), and other genes encoding germin-like proteins, ankyrin repeat/tetratricopeptide domain containing proteins, sugar and potassium transporters, cysteine protease, glycosyltransferases, fasciclin-like arabinogalactan proteins, and endosperm-specific proteins. Many of these candidate genes involved in stress response, hormone metabolism (mostly gibberellin and ethylene), yield, circadian rhythm, and flowering regulations also showed significant epistatic interactions among each other including various uncharacterized candidate genes (Figure S2, Table S4.2). Some interesting candidate genes such as *OsCIN1, OsSAP5*, and *OsTPR043* showed multiple epistatic interactions with AC and PC associated genes encoding proteins such as post-translational modification kinases, wax synthase isoform, putative bZIP protein (LOC_Os02g13810 – *OsHEI10/Human Enhancer of Invasion 10*), signaling G-protein (LOC_Os02g25870), and tonoplast monosaccharide transporter 2 (LOC_Os02g13560), among other unknown genes linked with both AC and PC. *OsHEI10* also showed numerous epistatic interactions with significant SNPs of genes encoding galactose oxidase beta-propeller (*OsFBK7*: LOC_Os02g30210), glycosyltransferase 2 (LOC_Os02g32750), poly ADP ribose polymerase 3 (*OsEnS-38*: LOC_Os02g32860), and MYB family transcription factor (LOC_Os02g30700).

### Distinct metabolome profiles of the pools of high AC and PC versus low AC and PC lines

Interestingly, amino acid pathways displayed significant variations between HAHP and LALP (Figure 2 and S4-5, Table S5). Additionally, pathways related to lipid biosynthesis, such as fatty acid and linoleic acid metabolism, demonstrated notable distinctions between these two groups (Figure 2D and Figure S3-4). To further support the claim of amino acid accumulation in HAHP, the line namely HAHP_101 and the recipient line Samba Mahsuri was considered as control were selected to test the protein yield. Intriguingly, the protein yield after alkali extraction was 15.99% for HAHP_101, whereas only 8.00% of protein was retrieved from Samba Mahsuri (Figure 2E). When comparing the colorimetric features of the protein concentrate, HAHP_101 has slightly lower L* values but slightly higher a* and b* values, indicating a less light appearance compared to Samba Mahsuri (Table S6). In addition, most EAAs were significantly higher in the HAHP group than in the Samba Mahsuri (Figure 2F). Notably, lysine exhibited the highest content among the amino acids in HAHP_101 sample.

**Figure 2.**
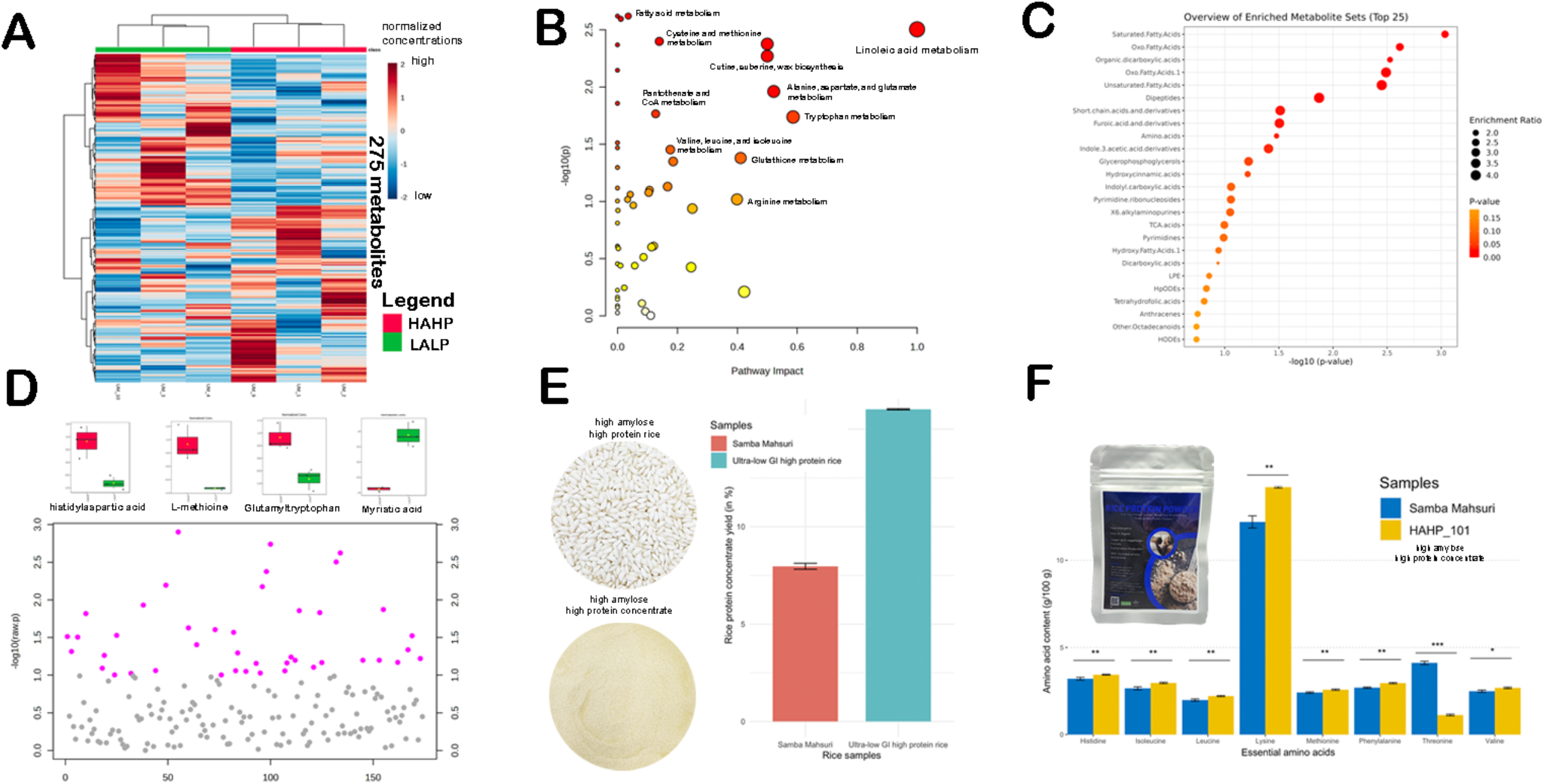
illustrates the metabolomic analysis of high amylose and high protein (HAHP) and low amylose and low protein (LALP) groups. (A) Heatmap distinguishing HAHP and LALP using 275 metabolites. (B) Pathway analysis indicating significant differences in amino acid and fatty acid-related pathways between HAHP and LALP. (C) Enrichment analysis highlighting the main class of compounds accumulating between HAHP and LALP. (D) T-test plot revealing the accumulation of amino acids and dipeptides in HAHP. (E) Isolated protein concentrate from the HAHP line showing a higher yield compared to Samba Mahsuri. (F) Rice protein powder from HAHP exhibiting higher levels of essential amino acids.

### Machine learning classification of ultra-low, low, and high glycemic index (GI) and their metabolomic signatures

A principal component analysis (Figure 3A) conducted on selected rice lines showed that GI classes (ultra-low, low, intermediate and high) showed clear distinction based on AC, with minor contributions from PC. Machine learning models using an artificial neural network developed following previous publications (7-9), optimized to 4-68-18-3 architecture also revealed that AC and PC in addition to two significant SNPs (snp_02_19362520 and snp_02_17959252) from *OsSBEIIb* and *OsFBK7* are key variables able to predict the GI class to an accuracy of 74.47%. Furthermore, the model showed robust predictive power for ultra-low GI and high GI lines, with true positive rates of 80.0% and 94.3%, respectively (Figure 3B), showcasing the high selectivity of the aforementioned GI classes. However, challenges arose in accurately predicting the low/intermediate GI class as individual genotypes belong to these classes may be misclassified as high GI (Figure 3B). Nevertheless, the machine learning approach demonstrated important phenome-genome parameters significantly influencing the GI trait distinguishing ultra-low GI from high GI categories within the RIL populations. The selected ultra-low GI and high protein lines are high yielding and early maturing compared to its to recipient parental line Samba Mahsuri (Figure 2F).

**Figure 3.**
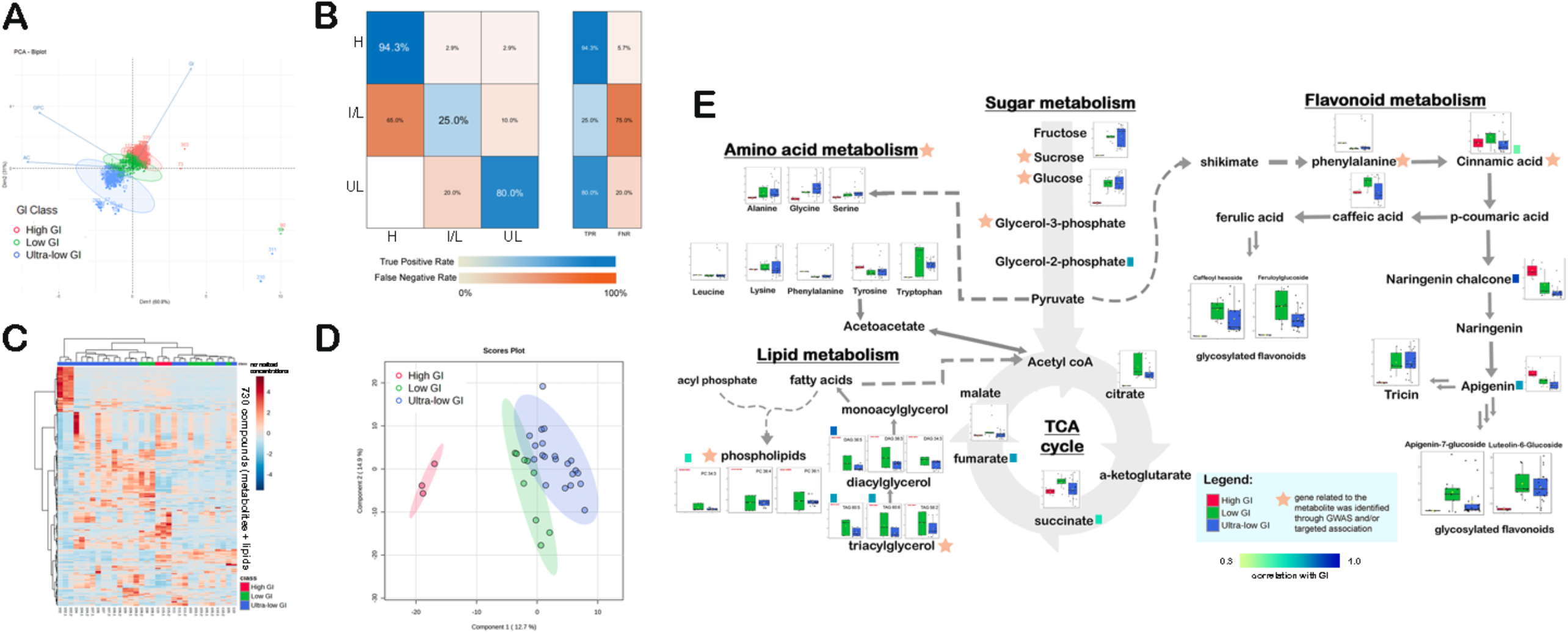
Modeling and metabolomic analysis of ultra-low, low, and high GI rice lines. (A) PCA showing the high, low, and ultra-low GI rice samples. (B) Confusion matrix demonstrating the accuracy of the artificial neural network (ANN) in classifying the ultra-low, low, and high GI rice lines. (C) Heatmap distinguishing the ultra-low, low, and high GI rice using the 730 metabolites. (D) PLS-DA model discriminating the ultra-low, low, and high GI rice samples using all the metabolites. (E) Summary pathway showing the differentially accumulated metabolites among the ultra-low, low, and high GI rice samples.

As illustrated in Figure 3C-D, the metabolites and lipids distinctly differentiate the low GI, ultra-low GI, and high GI lines (Table S7). Upon scrutinizing the primary metabolites from the GC-MS analysis of the rice samples, it became evident that various amino acid biosynthesis and sugar metabolism pathways differed between the ultra-low GI and high GI rice samples (Figure S5-S6). Consistent with this feature, the low and ultra-low GI rice samples exhibit higher levels of tryptophan and hydroxyproline. Furthermore, sugars show significant variation among the samples, with erythrose, glucuronic acid, maltose, and erythritol exhibiting the highest log-fold change, favoring the low and ultra-low GI rice samples. Besides amino acids, pathway analysis unveiled phenolic compounds such as tricin, caffeoylquinic acid, feruloylglucoside, *p*-coumaroyglucoside, luteolin-6-glucoside, hydroxygallic acid derivative, caffeoyl hexoside, and sinapoyl glucoside were found to significantly accumulate in the low and ultra-low GI rice lines (Figure S6). Conversely, the high GI line exhibited an accumulation of lipids classified as phosphatidylcholines, phosphatidylethanolamines, and their lyso-counterparts (Figure S7). Figure 3E provides a summary of the metabolite compounds mapped in their respective pathways, illustrating the preferential activation of these pathways. The preferential influx directed towards amino acid and flavonoid biosynthesis in ultra-low GI and HAHP lines and lipid accumulation in the high GI lines emphasize the potential significance of these pathways in contributing to the differences in glycemic index.

### CRISPR/Cas9-Mediated *SBEIIb* targeted mutagenesis in rice

An sgRNA was directed to induce mutations at the donor-splicing site of exon 15 of *SBEIIb (LOC_Os02g32660)* in the high yielding IRRI 154 line. T0 plants resulting from *Agrobacterium*-mediated transformation underwent PCR screening using pUbi-F3/Cas9-97-R primers to confirm T-DNA insertion. Subsequently, Cas9-positive plant DNA was utilized to amplify the target site with SBEIIb-TS1-F1/R1 primers (refer to Table S8). Twenty T0 individual events were then subjected to Sanger sequencing. Among the 20 T0 lines, nine mutant lines were identified comprising six heterozygous mutant lines, one biallelic mutant, and two lines exhibiting homozygous mutations. Mutation types included 1 bp insertion, 1 bp deletion, and large deletions up to 41 bp. Notably, line 35 was identified as homozygous with an ‘A’ bp insertion in both alleles, while line 46 exhibited ‘A’ and ‘G’ insertions on each allele. T1 seeds from lines #35 and #46 were utilized for *in-vitro* GI phenotyping and to estimate resistant starch content. These T1 seeds demonstrated reduced GI values of 55 and 54, respectively, compared to the wild type (GI of 59). Moreover, the percentage of resistant starch increased from 0.18% to 2.50% and 2.35% for lines 35 and 46, respectively (Figure 4). These two knockout lines, along with other mutant lines harboring different knockout mutations, will be grown for the T1 generation, and screened for transgene-free and homozygous lines through Mendelian segregation.

**Figure 4.**
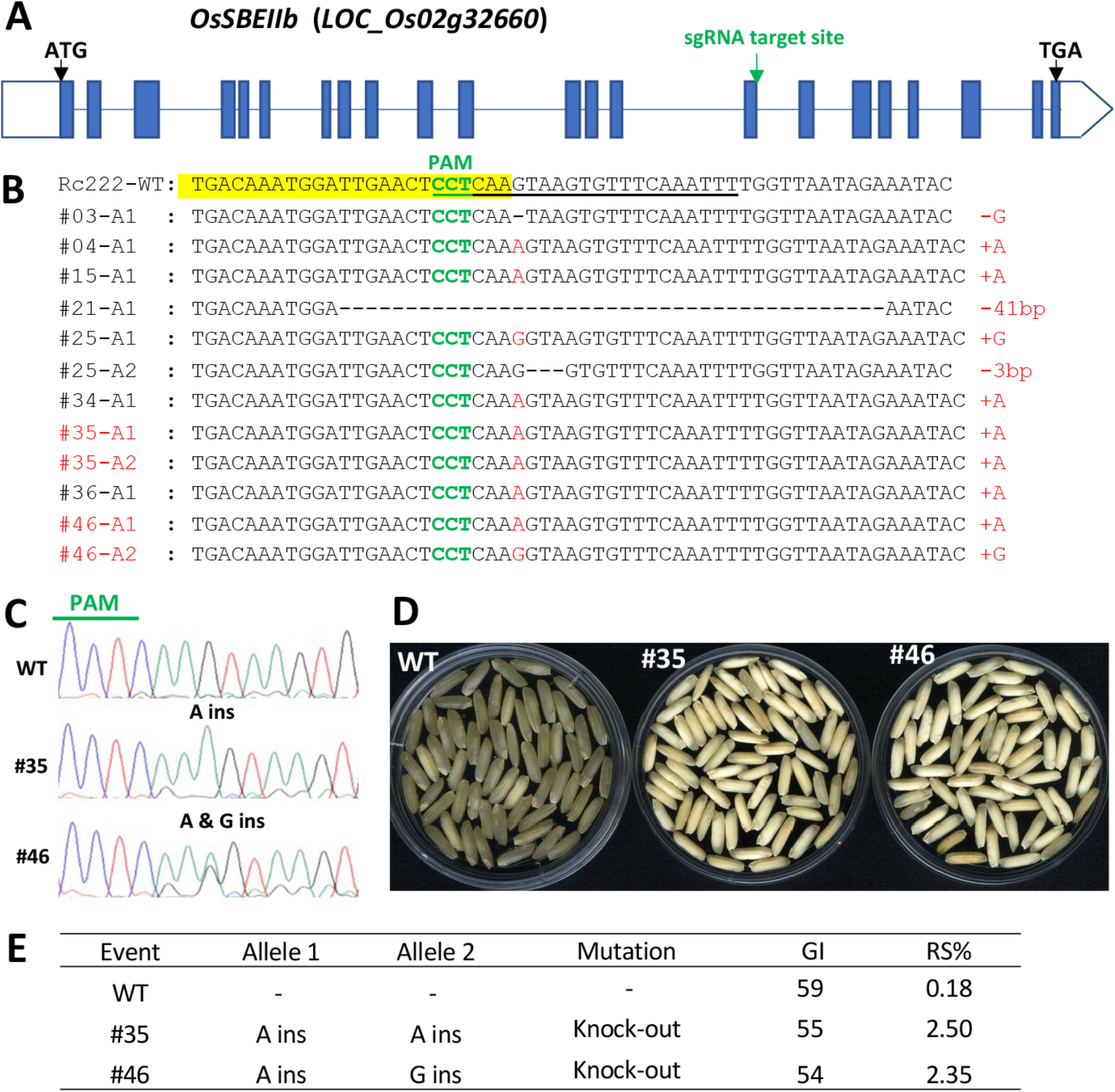
CRISPR/Cas9-mediated *OsSBEIIb* editing. (*A*) Schematic representation of *SBEIIb* gene with sgRNA target site. (*B*) Sequence comparisons at the CRISPR target site among WT (NSICRc222) and nine independent T_0_ plants. (Yellow highlight: 15^th^ exon region, underlined sequence: the sgRNA target site with PAM (protospacer-adjacent motif). (*C*) Sequencing chromatograms of the two selected knock-out (KO) T_0_ plants. (*D*) Morphology of the T_1_ seeds harvested from the two KO T_0_ plants. (*E*) Analyses of in-vitro GI and resistance starch (RS) with T_1_ seeds.

## Discussion

The nutrition quality of rice together with cooking and eating quality are largely determined by the composition of starch and protein in grain endosperms. Rice with low GI and higher protein content is a healthier option to support public health especially diabetic patients and for protein supplementation. In this study, we developed pyramided high amylose and high protein rice line exhibiting ultra-low glycemic index. This line was developed from the recombinant inbred lines from the parental lines of Samba Mahsuri (good grain quality) and IR36ae (high amylose & low GI) until the F_6_ generation. The present study implemented QTL-seq, QTL mapping, targeted association and epistatic interactions analyses and identified hotspot QTL and candidate genes associated with lowering GI while increasing the protein content. Metabolome analysis revealed differentially regulated metabolites and enriched pathways distinguishing the contrasting lines of HAHP and LALP having a range of glycemic index from ultra-low to intermediate classes which can be developed into functional food products.

### The genetics underlying low GI and high protein lines

Starch digestibility is affected by numerous extrinsic and intrinsic factors including genetic makeup, composition of starch and the extent of starch-protein interaction. High amylose crops are usually characterized with lower glycemic food load and higher resistant starch as opposed to low amylose lines which exhibit higher starch digestibility (10). Nonetheless, starch biosynthesis is a complex process regulated by multiple genes along with numerous transcription factors and mediators which further affect grain shape, weight, appearance, and overall quality. The QTL-seq analysis strategy considering whole-genome sequencing of pooled samples revealed numerous QTL regulating AC and PC, spanning several Mb regions. Fine mapping and validation of the QTLs was obtained by GBS of the F_5_-derived F_6_ recombinant inbred lines, composite interval mapping with targeted association analysis and epistatic interactions. Some significant genes identified in the hotspot QTLs (mostly ranging from less than 1Mb to 2Mb) in chromosome 2 included LOC_Os02g32660 (*SBEIIb*), LOC_Os02g15070 (*Glutelin B6/OsEnS-32*), LOC_Os02g33110 (*Cell-wall Invertase 1*), LOC_Os02g34560 (*OsNIN8/neutral/alkaline invertase 8*), LOC_Os02g31290 (*LARGE1* controlling grain size and weight), LOC_Os02g34630 (*MYB-related transcription factor*), three germin-like proteins, and seven ankyrin repeat/tetratricopeptide domain containing proteins.

By deploying high PVE contributed SNPs for low GI, high AC and high PC the modeling results were able to identify *sbeIIb* as an important locus. The G-to-A mutation at the splice junction between exon and intron 15 of *SBEIIb* leads to an early stop codon and results in a shorter SBEIIb underpinning low GI with chalky phenotype. Based on the QTL mapping results additional candidate genes influencing ultra-low GI phenotype with high protein were identified. Many studies have shown that inactivation of *OsSBEIIb* effectively increases the proportion of medium and long amylopectin chains (11) as well as resistant starch (12), and decreases levels of amylopectin chains (13), leading to overall lower starch digestibility. In the current study, we observed almost 60% variation in apparent amylose content and 8% variation in protein content attributed to a single nucleotide change in *<u>Os</u>SBEIIb*, further supporting previous knowledge on its substantial role in AC variation and demonstrated additional effects on the PC.

An important candidate gene among the additional ones we identified was the *OsCIN1*. This is involved in cleaving and unloading sucrose in the apoplast to the developing caryopsis during early grain filling stages. It plays complementary and synergistic roles with *OsCIN2/GIF1* and *OsCIN3* (14-16). Both *OsCIN1* and *OsCIN2* were reported to be extensively subjected to selection pressure over the years for better grain filling and potential yield (17). Another important candidate gene contributing to AC and PC is *OsNIN8* which encodes a neutral/alkaline invertase (Inv-N). These enzymes are known in catalyzing the hydrolysis of sucrose to produce glucose and fructose (18). Mutation of *OsNIN8* showed short root, delayed flowering, and partial sterility (19). Aside from *OsCIN1* and *OsNIN8*, multiple genes encoding glutelin, germin-like proteins, and ankyrin repeat/tetratricopeptide domain containing proteins were significantly detected in the current study. Germin-like proteins (GLPs) are characterized as water-soluble plant glycoproteins belonging to the cupin superfamily together with spherulins, vicilin-like and legumin-like seed storage globulins, and sucrose binding proteins (20). On the other hand, previous studies on some genes encoding ankyrin repeat/tetratricopeptide domain containing proteins (TPR) such as *FLOURY ENDOSPERM 2* were shown to positively regulate both starch biosynthesis and storage proteins biosynthesis (21, 22) by interacting with FLO2-interacting cupin domain protein 2 (FLOC1) and a bHLH transcription factor (23). Another tetratricopeptide-encoding gene, *OsCEO1*, was identified in a previous study as being a superior conductor of regulatory cascade of endosperm organogenesis controlling overall rice grain quality (24). Our current results suggest that several ankyrin/TPR-encoding genes might also be contributing to AC and PC variations, implicating the need for further functional investigations. We recommend doing in-depth experimental validation of the hotspot QTL regions detected in Chromosome 2 to elucidate the functions of additional candidate genes and their possible molecular interactions regulating ultra-low GI and high protein.

### Ultra-low GI and high protein lines share a preferential enrichment of glycolysis and amino acid biosynthesis

Homozygous *OsSBEIIb* mutant plants exhibiting high amylose in a previous study showed broad metabolic changes and transcriptional reprogramming with upregulation of genes encoding AGPase, soluble starch synthase and other starch branching enzymes, and downregulation of genes encoding granule-bound starch synthase, debranching enzymes, pullulanase, and starch phosphorylases (25). Similarly, we observed a range of differentially abundant metabolites in the recombinant inbred lines with different GI classification and high protein lines.

Amino acids usually serve as precursors for a wide array of both primary and secondary metabolites, act as intermediates in the production of end-product metabolites, and contribute to the regulation of multiple metabolic pathways (26, 27). The prevalence of amino acid pathways in the superior ultra-low GI group can be attributed to the close regulatory network between the aspartate family pathway and the synthesis of essential amino acids found in the samples. The high number of amino acids associated with the superior rice lines could indicate the effect of the level of free amino acids as precursors of protein accumulation in rice grains (28), since these ultra-low GI and low GI (superior lines) contain higher protein contents than the high GI ones. Moreover, dipeptides like glutamyltryptophan and histidylaspartic acid exhibited accumulations in the superior lines (Figure 3D). While the precise roles of these dipeptides remain undefined, it is possible that dipeptides derived from aromatic amino acids act as intermediates in the formation of flavonoids (27). In fact, Sadeghi et al. (29) reported that flavonoid glycosides inhibit α-glucosidase and α-amylase in both *in vitro* and *in vivo*. In addition, these flavonoids are known to ameliorate diabetes by regulating numerous cellular networks such as NF-κB, PI3K/Akt, MAPK, GSK3 and PPARγ. Both lysine degradation and biosynthesis were significant pathways associated with the high amylose/high protein group. High lysine content in endosperm, associated with *ae* and mutated *opaque* (*o2*) gene expression (30), has been correlated with an increase in starch biosynthesis enzymes (31). In contrast to the accumulation of intermediates from amino acid metabolism for fatty acid biosynthesis, our findings in the high amylose/high protein group reveal a redirection of flux towards glyoxylate/dicarboxylate metabolism. The glyoxylate metabolism primarily takes place in the peroxisomes and converts the acetyl-CoA, produced through the ß-oxidation of fatty acids, into succinate. Succinate is then utilized for carbohydrate synthesis (32). These results suggest that the glyoxylate cycle might play a role in the metabolic shift observed in superior ultra-low GI combined with high protein rice lines, where amino acid metabolism pathways dominate. In contrast, among inferior rice lines with high GI and intermediate protein, where the metabolic shift is geared toward fatty acid biosynthesis. The lipid-associated metabolic pathways in the low amylose group show the competition between carbohydrate metabolism and fatty acid synthesis (33). In particular, saturated fatty acids such as myristic acid and undecanedioic acid are accumulating in the LALP group. In general, fatty acid content in cereal grains influences lipid stability, and affects functionality properties during processing and storage (34). Vital metabolic changes for carbohydrates, protein, and fatty acids were corroborated with the findings identified in QTL-seq results in the present study. Candidate genes identified for exhibiting underlying variations could provide promising linkages to vital metabolic fluxes being regulated. This additionally warrants further functional validation for such candidates.

### Rice protein supplemented with enrichment of essential amino acids

Our current study reports a protein level of 15.99% in the ultra-low GI line, representing an almost 500% increase over conventional milled rice (brown rice ≈ 7.4 g, and white rice ≈ 2.7 g) (35). Additionally, the higher levels of essential amino acids (EAAs) found in the HAHP sample could significantly contribute to meeting the Recommended Dietary Allowance (RDA) for amino acids (31). For instance, 100 g of the HAHP rice provides EAAs such as isoleucine (2.96 ± 0.04 g), leucine (2.21 ± 0.03 g), lysine (14.19 ± 0.05 g), methionine (2.57 ± 0.04 g), phenylalanine (2.95 ± 0.04 g), and valine (2.67 ± 0.04 g), which can meet the RDA for individuals aged 10 and above (31). This preferential enrichment of essential amino acids with very high levels of protein may occur through the action of enzymes like aspartate kinase, argininosuccinate synthase, and aspartate semialdehyde (36). Notably, previous studies have demonstrated that the modulation of feedback inhibition of enzymes such as aspartate kinase and dihydrodipicolinate synthase led to a substantial increase, up to 21.7-fold, in lysine levels in rice through a transgenic strategy (37). Collectively, these findings underscore the potential of HAHP rice as a nutritional powerhouse, offering a substantial source of protein and essential amino acids that could play a pivotal role in meeting dietary requirements and addressing nutritional challenges, particularly in regions where rice is a dietary staple.

## Materials and Methods

Biochemical, genetic materials, and molecular procedures are described in SI Appendix, SI Materials and Methods.

## Data Availability

All study data are included in the article and SI Appendix.

## Author Contributions

SB: Conceived the QTL Seq strategy and conducted interpretation of QTL-seq data, performed breeding of RIL population, methodology; EAP: QTL analysis, targeted associations and epistatic interactions, writing - original draft; GM: QTL-seq analysis, methodology; VP: amino acid analysis; RNT: Metabolomics data analysis and interpretation, writing; RJB: Data analysis, modeling, writing; ARC: *in vitro* GI experiments, JS: Metabolomics data analysis and interpretation, SA: Metabolomics data analysis and supervision, AF: Metabolomics data analysis and supervision, AK: guidance and editing, NS and G.S.K: conceptualization, writing – editing and revisions, NS: acquisition of funds.

## Competing Interest Statement

The authors declare no conflict of interest.

## Acknowledgments

NS acknowledges funding from the Foundation for Food and Agricultural Research (CA-21-SS-0000000157), Department of Agriculture and Farmers Welfare, Government of India, The Indian Council of Agricultural Research (ICAR). We would like to acknowledge the Department of Science and Technology - Advanced Science and Technology Computing and Archiving Research Environment (DOST-ASTI COARE) for providing facilities for most of our analyses. We thank Drs. Tobias Kretzschmar and Juan David Arbelaez from IRRI for providing kind support for initial KASP maker development and conducting initial segregating studies, and Dr. Mallikarjuna Swamy from IRRI for providing his valuable suggestions on the breeding methodology adopted. We also thank Socorro Carandang for DNA extraction and acknowledge the support of genotyping by sequencing services (GBS) of The Elshire Group Limited. We thank CSIRO Australia for offering the in vitro GI measurement services and Integrated Service Laboratory (ISL) at IRRI for protein and amylose estimation of samples and cross-cutting service team for setting up crosses, population advancement and field support. We also like to thank Jazlyn Uy for isolating the rice proteins. R.N.T. acknowledges the Academy for International Agricultural Research (ACINAR) for funding his Ph.D. ACINAR, commissioned by the German Federal Ministry for Economic Cooperation and Development (BMZ), is being carried out by ATSAF (Council for Tropical and Subtropical Agricultural Research) e.V. on behalf of the Deutsche Gesellschaft für Internationale Zusammenarbeit (GIZ) GmbH. S.A. and A.R.F. acknowledge the European Union’s Horizon 2020 research and innovation programme, project PlantaSYST (SGA-CSA No. 739582 under FPA No. 664620) and the BG05M2OP001-1.003-001-C01 project, financed by the European Regional Development Fund through the Bulgarian ‘Science and Education for Smart Growth’ Operational Programme.

## Figures and Tables

**Table 1.**
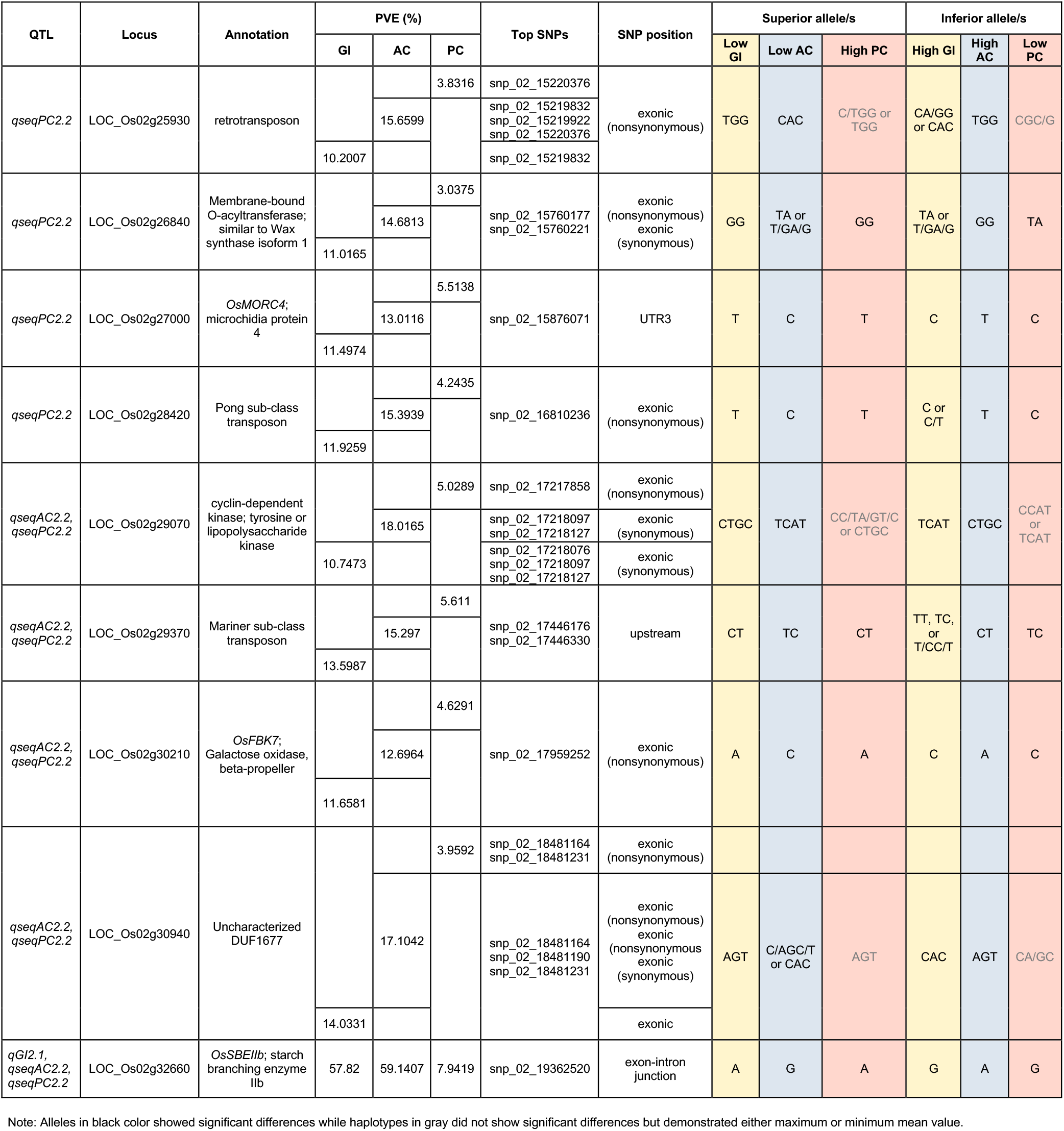
Candidate genes from significant QTLs based on BSA-seq analysis and targeted association analysis of recombinant inbred lines.

## Notes

### Competing Interest Statement

The authors have declared no competing interest.

