## Supplementary material for "Multi-omics of a rice population identifies genes and genomic regions in rice that bestow low glycemic index and high protein content": SI Appendix, SI Materials and Methods

**Development of a $F_2$-derived $F_3$ population.** The $F_2$-derived $F_3$ mapping population was developed by employing the hybridization between Samba Mahsuri (a superior grain quality line with dominant allele of *OsSBEIIb*) and IR36 amylose extender (IR36*ae*) line(recessive allele of *OsSBEIIb*), a high amylose line mutant created using the IR36 (1, 2). Samba Mahsuri, one of the parents, is one of the most popular varieties consumed mainly across India due to its excellent cooking and eating qualities. A total of ~5500 F2 progenies of IR36ae x Samba Mahsuri were raised. Upon examining the SNP variations existing in *SBEIIB* (LOC_Os02g32660) gene between both parents, one SNP (snp_02_19362520, G→A) previously reported by Butardo et al. (3), predicted to be located at the exon-intron junction (Exon 11 coordinates 19362521-19362640) was genotyped using Kompetitive allele specific PCR (KASP) assay. After assessing their fit to the Mendelian segregation ratio, a total of 369 $F_2$ progenies were selected and eventually self-pollinated until $F_3$ generation utilizing single seed descent (SSD) method and advanced to $F_6$ generation under open field conditions.

**Measurement of Amylose and Protein Content.** Seeds collected from $F_3$ generations of each family were bulk-harvested and dried until seed moisture content of 12-14% was attained. The samples were subjected to milling and the homogenized rice powder was subjected to amylose and protein estimation. The apparent amylose content (AC) was determined using an in-house validated method by the IRRI Service Laboratories (4) which was based on ISO 6647-2:2020 (Rice—Determination of Amylose Content—Part2: Routine Methods) (5) method. The apparent amylose content (AC) was determined using an in-house validated method by the IRRI Service Laboratories (4) which was based on ISO 6647-2:2020 (Rice—Determination of Amylose Content—Part2: Routine Methods) (5) method. The test solution is prepared by hot-dispersion with sodium hydroxide and analyzed using the San++ Segmented Flow Analyser (SFA) system (Skalar Analytical B.V., The Netherlands) and allowed to react with iodine to form amylose-iodine complex (K-I2). The absorbance of the iodine complex solution was determined at 620 nm wavelength and the AC (%) was quantified against the standard curves.

Crude protein content (PC) was measured using an in-house protocol of the IRRI Service Laboratories (54). A test portion of rice flour (50 mg) was digested in sulfuric acid (2 mL) with 1 g anhydrous potassium sulfate:selenium mixture (50:1, w/w) for 1 h at 380 °C. The sample was then cooled to room temperature, made up to volume (20 mL) with deionised water, and kept overnight to allow for sedimentation. The liberated ammonium in the digest was allowed to react with the solution consisting of sodium salicylate and sodium nitroprusside , and aqueous sodium hypochlorite through the glass transition lines in a Continuous Flow Analyser (CFA) system. The absorbance values of the ammonia-salicylate complex were determined at $\lambda = 660$ nm, and the % Kjeldahl N values of the samples were determined based on the non-linear relationship between absorbance and analyte concentration, following the Beer-Lambert Law based on a standard curve developed using ammonium sulfate solutions with different concentrations. Finally, the crude PC was calculated by multiplying the Kjeldahl N value by 5.95 (6).

***In-vitro* GI and resistance starch analysis.** The in vitro GI values of the milled rice samples of the $F_5$ derived $F_6$ RIL mapping populations was measured at the Commonwealth Scientific and Industrial Research Organisation (CSIRO), Adelaide, Australia using a predictive in vitro protocol that mimics the oral, gastric, pancreatic and intestinal digestion process in the human gut (7). The milled rice samples of $T_0$ homozygous lines of sbe2b genome edited lines and the corresponding wild type samples were subjected to the *in vitro* starch digestion was followed according to section 2.3.3 Method 3 described in Pautong et al. (8) with some modifications. Detailed method was described in the Supplementary information. In summary, the process involved cooking 300 ± 0.3 mg whole milled rice with a 1:2 rice-to-water ratio in a 50-ml tube over boiling water for 23 minutes. The sample was then left at room temperature for 5 minutes before it was transferred in a beaker fitted to the NutraScan GI20 Glycemic Index Analyser. It was added with 1mL of 0.05M sodium potassium phosphate buffer (pH 6.9) and the rice was minced using a stainless-steel spatula. The temperature was maintained at 37◦C. Subsequently, the 50-ml tube was washed with 4mL of the same buffer and the washing was also transferred to the beaker. To simulate the digestion process, the NutraScan GI20 Glycemic Index Analyser was used to dispense the necessary enzymes and buffers in the following manner: 2 mL of α-amylase in 0.05 M sodium potassium phosphate buffer (pH 6.9), 3 mL of aqueous HCl (pH ~1), then 3 mL of pepsin in 0.1 M sodium potassium phosphate buffer (pH 1.5). It was allowed to incubate for 30 minutes before the addition of 15 mL of aqueous NaOH (pH ~12.6). Following this, a 15 mL enzyme solution containing pancreatin and amyloglucosidase in 0.1 M sodium potassium phosphate buffer (pH 6.9) was added and was allowed to incubate for another 30 minutes. Duplicate aliquots (0.70 mL) were withdrawn at 0 minutes (just before pancreatin-AMG addition) and at 30 minutes (after pancreatin-AMG addition). The aliquots were placed on ice until centrifugation (13,500 rpm, 10 minutes, 4 ◦C). After which, a 1-μL aliquot of the supernatant was transferred to a 0.6-mL microfuge tube and added with 9 μL of AMG solution (33.3 U/mL in 0.4 M sodium acetate buffer, pH 5.0), vortexed, and incubated in a water bath (50 ◦C) for 20 minutes. Lastly, glucose in the mixture and reagent blank were measured by adding GOPOD reagent. The absorbance values at 510 nm for both the sample and glucose standard were corrected using a reagent blank. The corrected values were then converted into glucose concentrations (mg/ml of aliquot) following the equation described by Pautong et al.(8) in section 2.3.4 and the area under the curve (AUC) was calculated based on the trapezoidal rule. Finally, the predictive GI (pGI) of the samples were computed using the linear equation pGI = 0.07821 x AUC (corrected) + 47.8876.

**Bulk generation, DNA isolation and whole genome sequencing.** Based on the extreme phenotypes of grain quality phenotype including AC and PC, the 14-15 progenies per bulk were selected from either of the extreme phenotypes and bulk sets were created for each respective phenotype, constructing a total of four bulk sets. Bulk sets created for amylose content were confirmed for their contrasting glycaemic index values, i.e. high and low amylose bulk sets served as low and intermediate GI bulks, respectively. Genomic DNA was extracted from the selected bulk set individually using the cetyltrimethylammonium bromide (CTAB) method and purified using Qiagen DNeasy Plant Mini Kit. Qualitative analysis of DNA quality was done by running the samples on 1% agarose gel and by performing the DNA quantification through NanoDrop 2000. The final DNA concentration was adjusted to 100 ng/μL. The 30X- whole genome resequencing (WGS) was performed for all four bulk sets, aside from parental genotypes. Before library preparation, samples were ensured for the right concentration using fluorometer (Qubit Fluorometer, Invitrogen) and DNA sample integrity was assessed using agarose gel electrophoresis using 1% Agarose Gel (150 V for 40 min). 1μg genomic DNA was randomly fragmented, followed by purification and end repair performed using AxyPrep Mag PCR clean up Kit. The repair DNA were combined with A-tailing mix followed by incubation at 37℃ for 30 min while Illumina adapters were ligated to adenylate 3' ends of the DNA following incubation of 16℃ for 16h. Multiple rounds of PCR amplification were conducted to enrich the adapter-ligated DNA fragments. The 150 bp pair-end reads sequencing of qualified libraries was done using Illumina Hiseq X10 sequencer. The genotype data processing and QTL-seq analysis were detailed in Supplementary information.

**Genotype data processing and QTL-seq analysis.**

The clean reads from the extreme bulk sets for respective traits mapped onto the Nipponbare pseudomolecule reference (MSU version 7) and the variants were called. SNP calls with reference allele frequency of 0.2 in both bulks were filtered out and removed as these may be due to sequencing errors. Next Generation Sequencing (NGS)-based BSA analysis was performed following the earlier reports (9, 10). Briefly, SNP index was calculated as the ratio of the alternative allele reads to the total read depth in the individual bulk samples. Then, delta SNP index was calculated as per Takagi et al. (11). The genome wide G statistics (G value) was calculated for each SNP by observed and expected allele depth of the reference and alternate allele assuming that the allele depth is equal for both resistance and susceptible bulks. G' value was calculated using a tricube smoothed G value by average weighted of the physical distance across the neighbouring SNPs within a given window that accounts for Linkage disequilibrium (LD) and also minimizes background noise attributed to SNP calling errors (9, 10). Both of the parents, IR36ae and Samba Mahsuri were used as references while calculating the key parameters of G', delta SNP, and P values, and only peaks identified contrasting in both parents were used for analysis.

QTL Seq analysis or Bulk segregant (Bulk Seq) was performed on bulk sets created based on extreme values of amylose content and protein content. Bulk Seq analysis was implemented using the G statistics, wherein G' has been adopted avoiding the background noise during analysis while accounting for linkage disequilibrium (LD) between SNPs. Significant QTLs associated with AC and GPC were identified based on -log10P, G' and delta SNP index based on the default thresholds setting of the pipeline. The genotyping-by-sequencing data was generated for $F_5$-derived $F_6$ population following the Elshire et al. (12) method and have incorporated the following changes: 100 ng of genomic DNA were used, 3.6 ng of total adapters were used, the genomic DNAs were restricted with ApeKI enzyme, and the library was amplified with 18 PCR cycles. The 150-bp paired end reads were sequenced using the NovaSeq sequencing platform. The ABH format genotype data was created using PLINK1.9 (13) with 20% missingness call rate resulting in 44,946 SNPs after filtering.

**QTL mapping, targeted association and epistatic interaction analysis.** QTL mapping analysis for protein and AC was performed for the $F_5$-derived $F_6$ recombinant inbred line population using composite interval mapping (CIM) method from the R/qtl package (14). Targeted association analysis of the candidate genes lying within the fine-mapped QTLs identified from BSA-seq and CIM analyses was conducted using PLINK1.9 (13) and EMMAX (15) and significant SNPs were filtered at 95% confidence level. Epistatic interactions among significant SNPs were also tested using PLINK1.9 at $p<0.05$. Candidate genes common to AC and PC based on either BSA-seq analysis or CIM analysis, and genes with multiple epistatic interactions were used to construct the association network summary using Cytoscape (16). The percent phenotypic variation explained by the markers were calculated using LDAK (17). The expression profiles of candidate genes common to AC and PC based on BSA-seq and targeted association analyses were checked in the Rice eFP Browser (http://bar.utoronto.ca/) and genes with absolute expression value > 1000 in developing embryos were selected and their significant SNPs were used to determine different allelic combinations showing distinct GI, AC, and PC values in the $F_5$-derived $F_6$ RILs visualized using the *ggstatsplot* (18) R package.

**Sample Preparation and Extraction of Tissue Metabolites.** Brown rice flour samples of QTL-seq pools of high amylose and low amylose, and high protein and low proteins were added with 800 µL of 80% methanol and homogenized for 90s. The homogenized samples were sonicated for 30 minutes at $4^{O}C$. Samples were then kept at -40°C for 1h, vortex mixed for 30s and kept for another 30 min. This was followed by centrifugation at 12,000 rpm and 4°C for 15 mins. The supernatant was transferred to a vial for LC-MS analysis.

**Metabolomic analysis of ultra-low, low, and high GI samples.** The extraction and processing of the samples for metabolomic analyses were performed following the previous study (19, 20) . Briefly, 50 mg of the rice samples were extracted with 1.2 mL MTBE:methanol. The upper lipid-containing phase (500 µL) was dried in a SpeedVac concentrator and then resuspended in a 250 µL acetonitrile:2-propanol (7:3, v/v) solution. Then, a volume of 2 µL per sample was injected into a Waters Acquity ultra-performance LC system with an RP C8 column coupled with Fourier transform MS in positive ionization mode. The workflow included peak detection, retention time alignment, and removal of chemical noise. Identified lipids were confirmed through manual verification of the chromatograms using Xcalibur (version 3.0, Thermo-Fisher, Bremen, Germany). Mass spectra were acquired using an Orbitrap high-resolution mass spectrometer: Fourier-transform mass spectrometer (FT-MS) coupled with a linear ion trap (LTQ) Orbitrap XL (ThermoFisher Scientific, https://www.thermofisher.com). Chromatograms and mass spectra were evaluated using Chroma TOF 4.5 (Leco) and TagFinder 4.2 software. Metabolite data correlation was analyzed using Expressionist Analyst 14.0.5 (Genedata, Basel, Switzerland) (https://www.genedata.com/products/expressionist). The metabolite reporting checklist is given in Supplementary Table 5.

**Mathematical Modeling for GI Classification, and Metabolomic Multivariate Statistical Analysis.** To classify each rice line independently to three GI classes (Ultra-low, Low and Intermediate, and High), the peak SNPs, significant targeted association SNPs, and grain quality parameters were used as input traits through the models following the previous works on classification modeling in rice (21-23). The phenome-genome models were created using MATLAB® (version R2023b) using either Random Forest or Artificial Neural Network, whichever showed higher accuracy. The hyperparameters for both models were optimized using a built-in Bayesian Optimization option in the software. The data set (n = 386) was split into training and testing sets with a 70:30 training-testing split ratio. The models were validated using 10-fold cross-validation. The selected phenome-genome parameters were combined to further improve the accuracy of the model. The processed metabolomic datasets were imported into a web-based software package, MetaboAnalyst v5.0, for principal component analysis (PCA) and orthogonal partial least squares discriminant analysis (oPLS-DA) to visualize the differences in metabolite between the two groups (24). Variable Importance in Projection (VIP) analysis in the oPLS-DA model was used to identify those metabolites having the highest discrimination potential (VIP score ≥ 1). Pathway enrichment analysis was performed using the *O. sativa* japonica as the KEGG pathway reference (24).

**Purification of rice protein isolate and quantification of amino acids.** The protein concentrate was isolated from the rice samples using the alkali extraction method as described by de Souza et al. (25), with slight modifications. In brief, 300 g of rice samples were combined with 1500 mL of 0.18% NaOH and stirred continuously for 30 minutes. The sample was then centrifuged to collect the protein extract, and the pH was adjusted to 4.8 by adding 0.1 M HCl. The resulting protein concentrate was obtained, and after centrifugation to remove the supernatant, it was neutralized and dried to yield the protein powder. The amino acids were quantified using the method reported by Sekhar and Reddy (26) with slight modifications. For amino acid quantification, seeds from all varieties were ground into a fine powder for analysis. Approximately 0.5-1.0 grams of each sample were weighed and transferred to a hydrolysis tube, followed by the addition of 3 ml of 6 N HCl. The tubes were sealed and placed in an oven at 110 °C for 24 hours for hydrolysis. After hydrolysis, the samples were diluted to 15 ml with borate buffer to adjust the pH to 6.5-7.5. The resulting solution was kept in a water bath at 37°C for 1 hour, and then filtered through a 0.22 µm filter into a 1.5 ml HPLC vial for subsequent analysis. The vial was then prepared for analysis (26).

**Vector construction and rice transformation.** The single guide RNA (sgRNA) which specifically targets the 15$^{th}$ exon of *OsSBEIIb* was designed by using the CRISPR RGEN tools (http://www.rgenome.net/about/). The 20bp gRNA with 4bp-overhangings was prepared by oligomer duplex (Table S1). And then, the short DNA duplex was ligated into the CRISPR-Cas9 binary vector (pSR339) digested with *AarI* restriction enzyme. The pSR339 vector was developed at the IRRI for a PCR-free single step CRISPR vector construction and it contains three cassettes on its T-DNA region: *pZmUbi1-SpCas9-tNOS* for expressing Cas9 protein, *pOsU3*-AarI-*LacZ*-AarI-gRNA scaffold for expressing sgRNA, and *p35S-HPT-tCaMV* as plant selection marker. After insertion of the oligomer duplex into pSR339, the sequence of the construct was confirmed by Sanger sequencing with Tnos-F primer. The final construct was transformed into *Agrobacterium tumefaciens* LBA4404 strain by using freeze-thaw method. For the *OsSBEIIb* gene editing, a popular high-yielding *indica* variety (NSICRc222, IRRI designation-IRRI 154) was used as a background material. Rice transformation was conducted by using immature embryo based on an efficient *indica* rice transformation method (27). Through rice transformation, more than 80 $T_0$ plants were produced. Information of all the oligomers used in this study is presented in Table S1.

**Sequence analysis of the *OsSBEIIb* target site.** To confirm the sequence of the CRISPR target site, PCR and Sanger sequencing were performed. Briefly, DNAs were prepared from $T_0$ plants by using a simple DNA extraction method (28). PCRs were conducted with two PCR primer sets: pUbi-F3/Cas9-97-R primer set to check T-DNA presence and SBEIIb-TS-F1/R1 primer set to amplify the *OsSBEIIb* target site. All the $T_0$ plants generated possessed the T-DNA. Thus, the target site amplicon from randomly selected 10 $T_0$ plants were sequenced by AB3730 DNA sequencer through the Macrogen, Korea (https://dna.macrogen.com/main.do#). Chromatograms of the sequencing results were manually analyzed.
